# Phylogenetics and structural modeling of centralspindlin and Ect2 provide mechanistic insights into the emergence of Metazoa and multicellularity

**DOI:** 10.1101/2024.08.16.607330

**Authors:** Michael Glotzer

## Abstract

Obligate multicellularity evolved at least 5 times in eukaryotes, including at the origin of Metazoa ^1, 2^. The molecular events leading to the acquisition of multicellularity are not understood in any lineage. Here, I present an integrated analysis into the emergence of three regulators of cytokinesis and the Metazoan kingdom. Phylogenetic analysis and structural modeling indicate that three interacting essential regulators of cytokinesis, Kif23, Cyk4, and Ect2, are highly conserved across all Metazoa. These proteins cooperate to link the plane of cell division with the position of the spindle during anaphase and subsequently nucleate the assembly of stable intercellular bridges ^3-6^, structures prevalent in Metazoan germlines ^7^. The closest relatives of Metazoa, Choanoflagellata, encode Kif23 and Ect2 orthologs. In contrast, choanoflagellate species variably encode members of paralogous sets of proteins related to Cyk4, some of which are predicted to interact with Kif23. These findings, in conjunction with prior knowledge, suggests that the evolutionary refinement of these three cytokinetic regulators was a proximal prerequisite for the evolution of defining features of Metazoa.

## Introduction

The Metazoan kingdom emerged 650-850 million years ago ^8, 9^, giving rise to all animals with obligate multicellularity as one of its defining features. While obligate multicellularity could rely solely on cell-cell adhesion, aggregation-based mechanisms for multicellularity are susceptible to mixing with genetically unrelated cells ^10^. Alternatively, or in addition, multicellularity can emerge from a series of mitotic divisions followed by incomplete cytokineses resulting in a cluster of interconnected cells ^7, 11-14^. This latter mechanism intrinsically ensures that the functions of a cell cluster are linked to a single genome upon which natural selection can act.

Not only are metazoa multicellular, all 5 clades of extant metazoa are diploid organisms that reproduce via eggs and sperm (anisogamy) ^7, 15-20^. Upon maturation/fertilization, oocytes undergo two sequential, highly asymmetric cleavages resulting in the extrusion of two polar bodies, generating haploid eggs. Thus, by inference, the last common ancestor of Metazoa (LCAM) had the ability to produce haploid sperm and eggs as well as mechanisms to cleave fertilized, diploid eggs into smaller cells that self-assemble into fertile organisms ^21, 22^. The cytokinetic machinery is directly involved in these processes.

Many of the proteins that promote cytokinesis in Metazoa are evolutionarily ancient ^7, 23^. Cleavage furrow ingression in cells from diverse species of Opisthokonts (Metazoa, Fungi, Amoebozoa, etc.) is driven by an actomyosin-based contractile ring (Figure 1A). Ring assembly in many opisthokonts is triggered by activation of the small GTPase Rho1 through nucleotide exchange ^23, 24^. Cytokinesis in metazoa requires three additional proteins: Ect2, Kif23, and Cyk4. Ect2 is the guanine nucleotide exchange factor (GEF) that activates Rho1 during cytokinesis (Figure 1A) ^25-27^, and Ect2 is activated by centralspindlin, a heterotetrameric complex that assembles from dimers of the kinesin-6 family member, Kif23, and Cyk4 (Figures 1A, B) ^28-34^.

**Figure 1:**
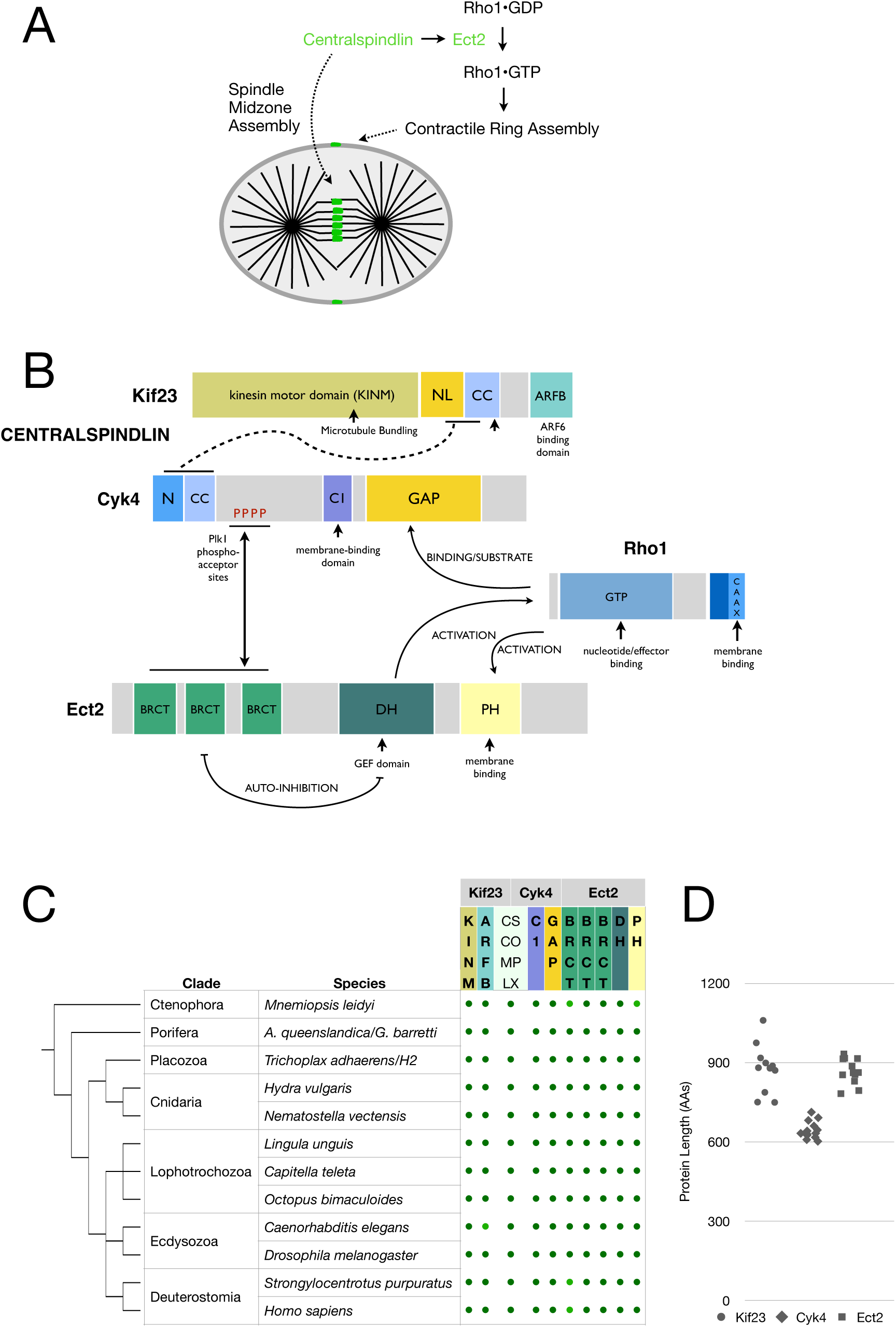
Centralspindlin and Ect2 are structurally conserved in all clades of Metazoa and regulate contractile ring and spindle midzone assembly. A. Schematic depicting the roles of Centralspindlin and Ect2 in cytokinesis in Metazoa. B. Detailed map of the known interactions between Centralspindlin subunits (Kif23/Cyk4), Ect2, and Rho1 (see text for details). C. Presence (green dot) of each structural domain (see B for domain abbreviations) in the Metazoan representatives of Kif23, Cyk4, and Ect2; a phylogenetic tree is provide for reference ^117^. A lighter green dot indicates lower confidence (< 70 pLDDT; predicted local distance difference test) in the structural prediction. The integrity of each domain was assessed by a combination of SMART, CDD-NCBI and Alphafold3. Centralspindlin complex formation (CS COMPLX) was assessed with Alphafold3 (see figure 2). D. Number of amino acids in Kif23, Cyk4, and Ect2 in species spanning metazoa listed in C.

Despite their comparatively recent evolutionary emergence, centralspindlin and Ect2 regulate many steps during cytokinesis. Association of centralspindlin with the anaphase spindle midzone promotes membrane accumulation of the centralspindlin/Ect2 complex at the cortical region overlying the mid-plane of the spindle, where it promotes local Rho1 activation, contractile ring assembly, and sustains ingression of the cleavage furrow (Figures 1A, B) ^3, 27, 35^. Following ingression, centralspindlin nucleates an intercellular bridge that physically connects daughter cells ^4-6, 36^. Many, but not all, cells eventually sever this bridge and undergo abscission, generating separate daughter cells ^37-41^. Thus, centralspindlin and Ect2 regulate each stage of cytokinesis in animal cells. Given the potential role of incomplete cytokinesis in clonal multicellularity, analysis of the evolutionary origin of these cytokinetic regulators could provide insight into the origin of multicellularity, a key feature of Metazoa.

## Results

### Centralspindlin and Ect2 are highly conserved in Metazoa

Orthologs of Kif23, Cyk4, and Ect2 are readily identifiable in representatives of all five clades of Metazoa. The domain structure and length of these three proteins is highly conserved among extant Metazoa (Figure 1C,D). Kif23 is a kinesin-6 motor protein paralogous to Kif20 (Supplementary Figure 1), a kinesin which contains an extended C-terminal coiled coil domain. Kif23 family members uniquely contain a C-terminal ARF6-binding domain (Figure 1B-C, Supplementary Figure 2) which associates with active Arf6 ^42, 43^. This interaction likely occurs on a membrane surface as Arf6 is myristoylated and membrane-associated ^44, 45^. Cyk4 contains a N-terminal coiled coil region and a membrane-binding C1 domain that is closely apposed to a RhoGAP domain ^36^ (Figure 1B). The Rho1 activator, ECT2, contains autoinhibitory tandem BRCT domains ^46^ (Figure 1B), a Dbl Homology (DH) RhoGEF domain, and a Pleckstrin Homology (PH) domain. The BRCT domains of Ect2 associate with Plk1-phosphorylated Cyk4 ^47-50^, promoting relief from autoinhibition (Figure 1B). Structure predictions ^51^ support the presence of three BRCT domains in representatives of Ect2 that span Metazoa (Figure 1C).

The interaction between Kif23 and Cyk4 is essential for centralspindlin function ^29, 52^. This interaction is mediated by the inter-digitation of coiled coil dimers of both Cyk4 and Kif23 along with their N-terminal flanking residues, forming an antiparallel helical bundle ^53-55^. The sequences of the interacting regions exhibit a discontinuous pattern of conservation (Supplementary Figures 2, 4). The N-terminal interaction domains of metazoan Cyk4 are predicted to form parallel coiled coils with flexible N-termini (Figure 2). To ascertain whether the structure of the centralspindlin complex is conserved throughout Metazoa, predicted structures of the corresponding regions of dimers of Kif23 and Cyk4 from a broad range of metazoa were generated ^56^. Cyk4 and Kif23 from species covering all metazoan phyla folded into similar tetrameric complexes (Figure 2, Supplementary Figure 5). These predicted folds closely resemble the experimentally determined structure of the corresponding region of *C. elegans* centralspindlin ^54^.

**Figure 2:**
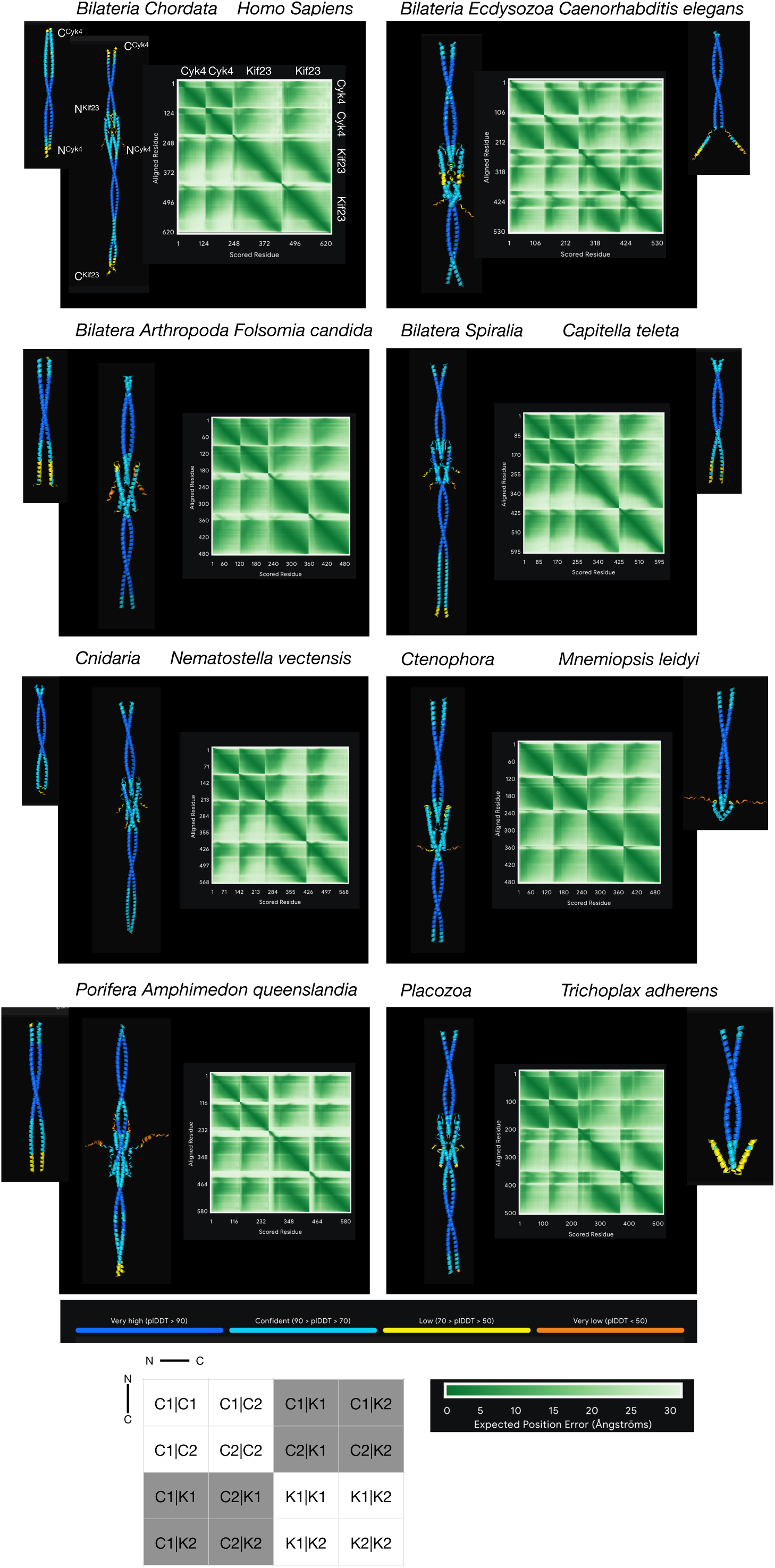
The interaction between Cyk4 and Kif23 is conserved among Metazoa. For representative species spanning Metazoa, the region of Cyk4 from the N-terminus to the end of the coiled coil and the region of Kif23 from∼30 amino acids from the beginning of the coiled coil to its end were used to generate Alphafold3 predictions of potential hezterotetramers; predictions were also generated for the Cyk4 homodimers alone (flanking heterotetramers). In the heterotetramers, Cyk4 subunits are positioned on the top and the N-termini of both protein segments are centrally positioned. Unstructured regions were trimmed. Confidence in structural predictions (pLDDT) of a given protein chain are color coded (see legend). Heterotetramer complex assembly is indicated in the predicted aligned errors (PAE) 2D plots; a low value (dark green) in the 2D plot indicates that the relative positions of a pair of residues are predicted with high confidence. Each sector in the PAE plots reflects the interaction between two chains (C:Cyk4, K:Kif23) as indicated in the table (unshaded sectors/ intradimer; shaded sectors/inter-subunit).

### Identification of proteins related to Kif23, Cyk4, and Ect2 in relatives of Metazoa

To determine the evolutionary history of Kif23, Cyk4, and Ect2, the taxonomic relatives of Metazoa were searched for related proteins. The next closest taxonomic clade to Metazoa is Choanoflagellata, followed by Filasterea and Teretosporea, collectively forming the Holozoa (Figure 3A) ^12, 57, 58^. The proteomes of representatives of these clades were analyzed for the presence of proteins with similarity to Kif20, Kif23, Cyk4, and Ect2. Orthologs of Kif20 and Kif23 were unambiguously identified in numerous *Holozoan* species (Supplementary Figure 1). Regions similar to the Arf6 binding domain are uniquely present in Kif23 family members, though the confidence of the structural predictions of these ARF6 binding domains vary considerably.

**Figure 3:**
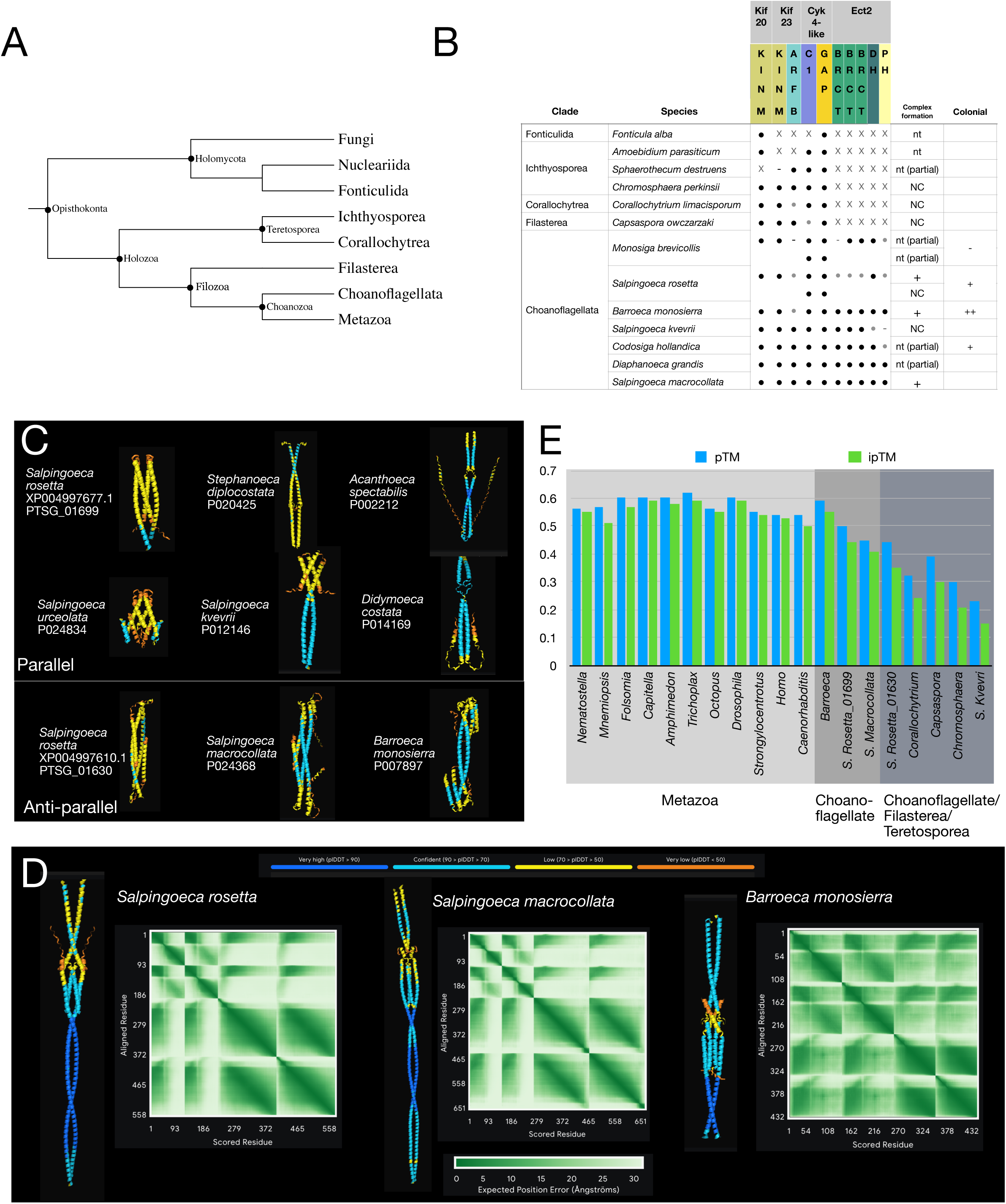
Distribution of Kif23, Cyk4, and Ect2 in Opisthokonta and the structure of putative centralspindlin-like complexes. (A) Phylogenetic tree of selected clades with Opisthokonta ^118^. (B) Presence or absence of proteins related Kif20, Kif23, Cyk4, and Ect2 and presence of domains in representative species. Black dot indicates presence of a given domain; gray dot domain is present, though structural predictions are low confidence (pLDDT < 70); X indicates not detected; - indicates domain status could not be determined as predicted proteins were incomplete. Complex formation is also indicated where + indicates ipTM > 0.4; NC (no complex) indicates ipTM < 0.4; nt indicates not tested due to absence of one or more proteins. Multicellular colony formation is indicated ^60, 62^. (C) Alphafold 3 predictions of the indicated N-termini of Cyk4-related proteins from choanoflagellates. (D) Alphafold 3 predictions of the heterotetramers of Kif23 and Cyk4-related proteins (S. Rosetta: PTSG06199/PTSG00966; S. macrocollata P024368/P024644; B. monosierra P000323/P007897); see figure 2 legend for a guide to the plots. (E) Summary statistics of the overall confidence of the Kif23-Cyk4/Cyk4-related heterotetrameric complexes, (see methods for details and Figures 2, 3, and Supplementary Figure 6 for predicted structures).

Unlike the clear orthology of Kif23 motor proteins amongst numerous holozoans, the C1:RhoGAP module characteristic of Cyk4 is also present in a diverse set of proteins in holozoans. Cluster analysis reveals that this set of proteins form nine subgroups (Supplementary Figure 3) with variable domains in their N-terminal regions: SH2, LIM, PH, F-Bar, myosin motor or a putative coiled coil, suggesting domain shuffling. Choanoflagellate proteomes contain numerous such proteins with C1:RhoGAP domains; the majority of such proteins contain N-terminal coiled coils. Nine choanoflagellate species contain two such paralogs (Supplementary Figure 3). Dimers of the N-terminal regions of these proteins are predicted, with low confidence, to form parallel or antiparallel coiled coils, as well as mixed helical configurations (Figure 3). Thus, the N-termini of choanoflagellate Cyk4-related proteins are structurally distinct from the N-termini of Cyk4.

Where permitted by sequence availability, structural predictions were generated for heterotetrameric complexes between the N-termini of choanoflagellate Cyk4-related proteins and the central region of their cognate Kif23 orthologs. In each case, the Kif23 regions are predicted to assemble into parallel coiled coil dimers, as is typical for plus-end directed kinesins (not shown). In three of five cases, the Cyk4-related proteins are predicted to form anti-parallel dimers that interdigitate with the cognate Kif23. The structures of these complexes are each somewhat distinct, consistent with the low sequence conservation of the interacting regions in both subunits (Figure 3, Supplementary Figures 2, 4). The parallel structure of the Kif23 dimer is predicted to induce two of the Choanoflagellate Cyk4-like proteins to adopt a parallel configuration, rather than the anti-parallel configuration predicted for the independent dimers. The predicted Kif23/Cyk4-related complex from *B. monosierra* is more similar to its metazoan counterparts, and the confidence score of the complex prediction is comparable to that of metazoan complexes (Figure 3D-E, Supplementary Figure 5). A further difference between choanoflagellate and metazoan complexes is apparent from the predicted structures: the surfaces of metazoan centralspindlin exhibit alternating basic and acidic patches, whereas the charged patches on the surfaces of the choanoflagellate complexes are predominantly acidic (Supplementary Figure 5).

Ect2 orthologs, as defined by the tandem BRCT:DHPH structure, were found in several choanoflagellate species, but not in clades more distant from metazoa. It remains to be determined whether choanoflagellate Ect2 proteins regulate cytokinesis and, if so, how they are activated; the conserved cluster of Plk1 phosphosites in metazoan Cyk4 are not conserved in choanoflagellate Cyk4-related proteins (Supplementary Figure 4).

The unicellular amoeboid Filasterean, *Capsaspora owczarzaki,* ^59^ contains a Kif23 ortholog. This organism also contains orthologs of the C1:RhoGAP domain proteins β-chimaerin, myosin IX, and GEM interacting protein. *C.o.* encodes a fourth C1:RhoGAP protein with a coiled coil region (Supplementary Figure 3). This protein, CAOG0002, clusters with proteins found in Teretosporea, is ∼2x longer than metazoan Cyk4, its C1 and RhoGAP domains are non-adjacent (separated by 173 residues), and the coiled coil is far (350 residues) from the N-terminus. Structural prediction of possible complex formation between the coiled coil region of this protein and *C.o.* Kif23 do not indicate that they interact (Supplementary Figure 6). No proteins containing both BRCT and RhoGEF domains could be identified in *C. owczarzaki*.

Similarly, *Teretosporea* encode Kif23 orthologs, but no proteins similar to Ect2 could be identified in this clade. While these organisms contain C1:RhoGAP domain containing proteins, they are otherwise unrelated to Cyk4 at the domain and sequence level and structural predictions do not provide evidence for complex formation with Kif23 (Supplementary Figure 3, 6).

## Discussion

Phylogenetic analysis reveals that Cyk4, Kif23, and Ect2 are highly conserved among all clades of Metazoa. Sequence conservation and structural modeling reveal that this conservation includes all known functional regions, including those that mediate assembly of the centralspindlin complex. While several non-metazoans contain Kif23 orthologs, only the closest relatives of metazoa, choanoflagellates, contain orthologs or paralogs of Ect2 and Cyk4. Importantly, choanoflagellates do not encode orthologs of Cyk4, but they do encode multiple paralogs. Despite the significant divergence from metazoan paralogs, some examples of complex formation between choanoflagellate Cyk4-related proteins with Kif23 is predicted, albeit with interaction interfaces that are distinct from those that mediate assembly of metazoan centralspindlin. These findings reveal that centralspindlin emerged contemporaneously with Metazoa, Ect2 emerged contemporaneously with Choanozoa, and these proteins are highly conserved in extant Metazoa.

Although choanoflagellates are typically unicellular, a number of species exhibit colonial behavior mediated by incomplete cytokinesis ^60^. Intriguingly, two of the species encoding putative centralspindlin-like complexes form multicellular clusters via incomplete cytokinesis: *S. rosetta* forms clusters in response to a bacterial signal ^61^ and *B. monosierra* forms large hollow spheres of bridge-linked cells containing bacteria ^62^. The striking structural similarities of the heterotetrameric *B.m.* complex to metazoan centralspindlin suggests convergent evolution of both structure and function. Finally, though Ect2 orthologs are present in choanoflagellates, the phospho-motif of metazoan Cyk4 that mediates Ect2 association is not evident in the choanoflagellate paralogs, indicating that Ect2 is likely activated by a different mechanism.

The conservation of Kif23, extending to *Teretosporea* and *Filasterea,* suggests that Kif23 may have first functioned independently of Cyk4-related proteins. By acquiring the ARF6 binding domain, Kif23 may have linked microtubule bundles in the spindle to membranes during anaphase.

### Centralspindlin and Ect2 are directly involved in functions core to Metazoan reproduction

What functional consequences might accrue to organisms from the acquisition of the cytokinetic regulators Kif23, Cyk4, and Ect2? The conservation of domain organization and protein length of Kif23, Cyk4, and Ect2 throughout metazoa suggest that, to a first approximation, observations made on extant centralspindlin/Ect2 are a reasonable proxy for the functions of these components in LCAM. Like metazoa, many opisthokonts, including some fungi, use a Rho-regulated contractile array of actomyosin to perform cytokinesis ^63^, suggesting that the ancestors of Metazoa were, unsurprisingly, cytokinesis competent. Thus, rather than enabling cytokinesis per se, acquisition of these proteins could have enabled novel functions. Prior analyses of centralspindlin/Ect2 have directly implicated these proteins in division plane positioning, asymmetric cell division, spindle midzone stabilization, cytokinetic bridge assembly, and cell fate specification and indirectly in adhesion between sister cells.

These processes are core to the mechanism of metazoan reproduction and multicellularity. The mechanism of division plane specification is not conserved amongst all opisthokonts. For example, the division plane of the most studied fungi, budding and fission yeast, are positioned prior to metaphase, through distinct mechanisms ^64, 65^. Likewise, division plane orientation in choanoflagellates appears highly stereotyped, forming between the flagella of the sister cells ^13, 14, 66^. In contrast, centralspindlin and Ect2 are directly implicated in partitioning daughter cells at a position dictated by the mid-plane of the anaphase spindle (Figure 4) ^26-29, 35, 67-69^. The linkage of the division plane to the spindle midzone not only prevents aneuploidy, but it also enables regulated spindle displacements to result in asymmetric cell division (Figure 4) and enables cells of vastly different sizes to be cleaved, as the mitotic spindle scales with cell size ^70, 71^. Indeed, an extremely asymmetric division occurs upon polar body extrusion during meiosis, a process that can involve centralspindlin (Figure 4) ^28, 72^. The egg, the large daughter cell of these highly asymmetric meiotic divisions, subsequently cleaves into progressively smaller blastomeres, reflecting the adaptability of centralspindlin-dependent cell division. Some blastomere divisions are asymmetric, generating daughter cells with distinct contents that alter their fates ^73, 74^.

**Figure 4:**
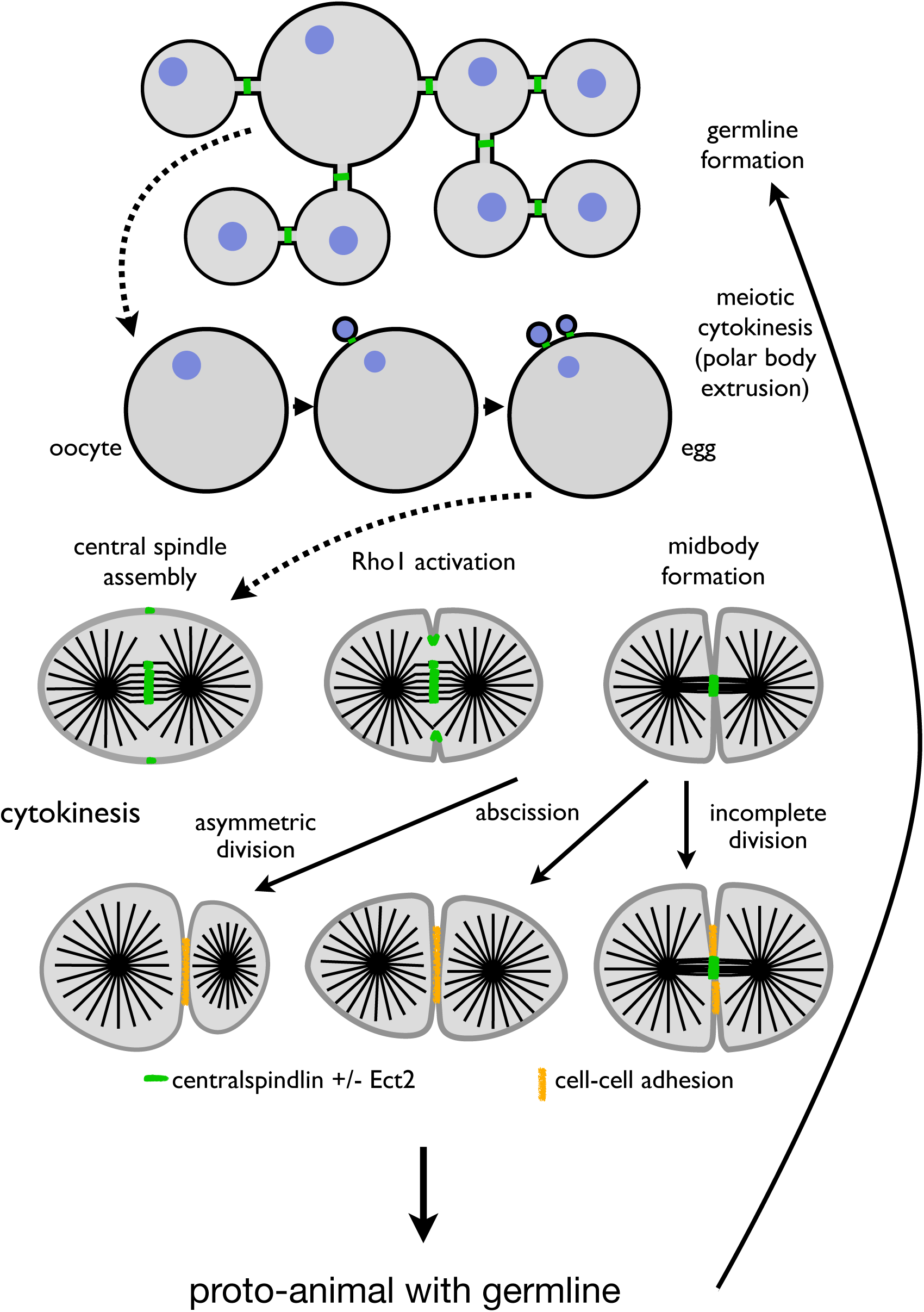
Centralspindlin enables keystone features of Metazoan reproduction. Schematic summarizing the roles of centralspindlin in metazoan reproduction and development. These include: stabilization of intercellular bridges to during germline development, positioning the division plane and activating Rho1 during meiotic and mitotic cell divisions (in conjunction with Ect2), promoting adhesion between sister cells during cell division, stabilizing persistent cytokinetic bridges, and enabling asymmetric spindle positioning to result in asymmetric cell division.

The events at the completion of cytokinesis substantively differ in Metazoa from those that take place in other Opisthokont species. Specifically, as furrowing nears completion, centralspindlin, along with Ect2, accumulates at the center of cytokinetic bridges^26, 28, 68^. This accumulation of centralspindlin bundles the associated microtubules ^69^ and stabilizes their association with the plasma membrane ^36^, forming a structure known as the Flemming body, or midbody. Ultimately centralspindlin reorganizes into a dense, membrane associated, ring at the center of the cytokinetic bridge. A plethora of proteins, not all of which are conserved throughout Metazoa, regulate abscission ^7, 75^. While many cells complete abscission, in certain cell types, cytokinetic bridges can undergo further maturation and stabilization ^76-79^, generating persistent intercellular bridges.

Intercellular bridges constitute a physical connection between related cells, generating a multicellular cluster. In metazoa, such connections are prevalent in, but not restricted to, male and female germline tissues, and, where studied, they involve centralspindlin (Figure 4) ^80-84^. In *Drosophila* female germlines, centralspindlin templates assembly of ring canals through which materials produced in the nurse cells are delivered into the oocyte as it develops ^85, 86^. Functionally analogous structures are present in nematodes and *Hydra* ^4, 87^ and contribute to fertility ^28, 87-89^. Cytokinetic bridges contribute to provisioning zygotes with the materials necessary for early development, until the nascent animal can acquire nutrients from its environment. Critically, such bridges are a conserved feature of germlines across Metazoa ^7^, suggesting they existed in LCAM.

Intercellular bridges also regulate animal development independent of cell division. Intercellular trafficking via bridge microtubule regulates cell fate decisions in mouse blastomeres ^90^. In nematodes, centralspindlin promotes formation of the pharyngeal epithelium ^91^. The Notch pathway, which also emerged at the onset of metazoa ^92^, can drive binary cell fate decisions, which, in some cases, depends on the spindle midzone ^93^. Cytokinetic furrowing and prolonged physical association between daughter cells as a result of delayed abscission can promote cell adhesion between daughter cells ^90, 94-97, 97, 98^, contributing to multicellularity. More broadly, cell lineages can play a central role in the establishment of developmental complexity^99, 100^.

Conceptually, the emergence of three interacting cytokinetic regulators, Kif23, Cyk4, and Ect2, paired a highly flexible mode of cytoplasmic division with the ability to control the length of time that daughter cells remain physically connected. The ability to prevent completion of cytokinesis itself generates a form of multicellularity, a mechanism intrinsically limits the cell cluster to cells of a common lineage, thereby protecting against genetically distinct cells ^13^. This protection would be particularly relevant in the germline, perhaps reflecting the remarkable conservation of intercellular bridges at that site ^7^. Taken together, the emergence of centralspindlin/Ect2 appears to have been directly involved in the contemporaneous acquisition of several key aspects of Metazoa: multicellularity, cell size heterogeneity, and anisogametic germlines (Figure 4). Analogous innovations that stabilize microtubules of the telophase spindle may have contributed to the emergence of multicellularity more than once during eukaryote evolution ^101, 102^.

## Supporting information

Sequence files

## Acknowledgements

This work was supported by NIH MIRA award 5R35GM127091. MG thanks his lab members past and present and Heather Marlow, David Pincus, Noah Mitchell, and Carlos Cortez (University of Chicago), Marc Kirschner (Harvard Medical School), Pierre Gönczy (ETH, Lausanne), Giacomo Glotzer (Rockefeller University), Ashley Rich (Duke University) and Nick Brown (University of Cambridge) for helpful discussions.

## Methods

### Bioinformatics

Orthologs of Kif23, Cyk4, and Ect2 were identified from the following databases: Uniprot ^103^, NCBI ^104^, and EukProt ^105^ with a primary focus on the Holozoan datasets ^58, 106-108^. In each case, Blast searches using the well conserved domains (Kif20/Kif23 motor domain, ARF6 binding domain, C1:RhoGAP, BRCT:DH:PH) were used to isolate potential orthologs. Initial hits were used for subsequent Blast searches in related species to identify further potential orthologs. The domain structures of these proteins were analyzed using a combination of Conserved Domain Database (CDD)^109^, SMART ^110^, Interpro ^111^, and Alphafold Structural Database and de novo predictions with Alphafold3 ^56, 112^. This approach readily identified Kif20/ Kif23 orthologs in Holozoa and Fonticulida; Ect2 orthologs in Choanoflagellata; and a large set of C1:RhoGAP domain containing proteins. To search for Ect2 orthologs in Filasterea, Icthyosporea, Corallochytrea, the available species from these clades in EukProt were searched with Metazoan and Choanoflagellate Ect2 orthologs. While proteins with DH or BRCT domains were readily identified, none of the identified proteins contained both DH and BRCT domains. Clustal ^113^ was used to generate multiple sequence alignments and phylogenetic trees using the motor domains of Kif23/Kif20 orthologs and the C1:RhoGAP domains of Cyk4-related proteins. The resulting alignments were visualized using WebLogo ^114^ and trees were generated with iTOL ^115^.

### Centralspindlin complex formation predictions

For each set of putative centralspindlin complexes, the corresponding orthologs of Kif23 and Cyk4 were analyzed with PairCoil2 ^116^ and the region of putative Cyk4 orthologs from the N-terminus to the end of the coiled coil was paired with the Kif23 sequences ∼50 amino acids N-terminus through the end of the coiled coil. These sequences were submitted to Alphafold3 specifying each protein was present in two copies ^56^. Predictions were refined by trimming the predicted unstructured termini. Formation of a complex was scored as positive if all the following criteria were met: (i) the structure exhibited antiparallel interdigitated helices, (ii) most of the interacting regions of the proteins scored > 70 in the predicted local distance difference test (pLDDT), (iii) extensive regions of the predicted align errors (PAE) plot that pertain to intersubunit interactions are <10, and (iv) the predicted template modeling (pTM) and the interface predicted template modeling (ipTM) scores were ≥0.4 ^56^. Although this threshold for pTM and ipTM is relatively low, these scores rarely exceed 0.6 even in the case of metazoan centralspindlin complexes, one of which matches the experimentally determined structure ^54^.

---

**Supplementary Figure 1:**
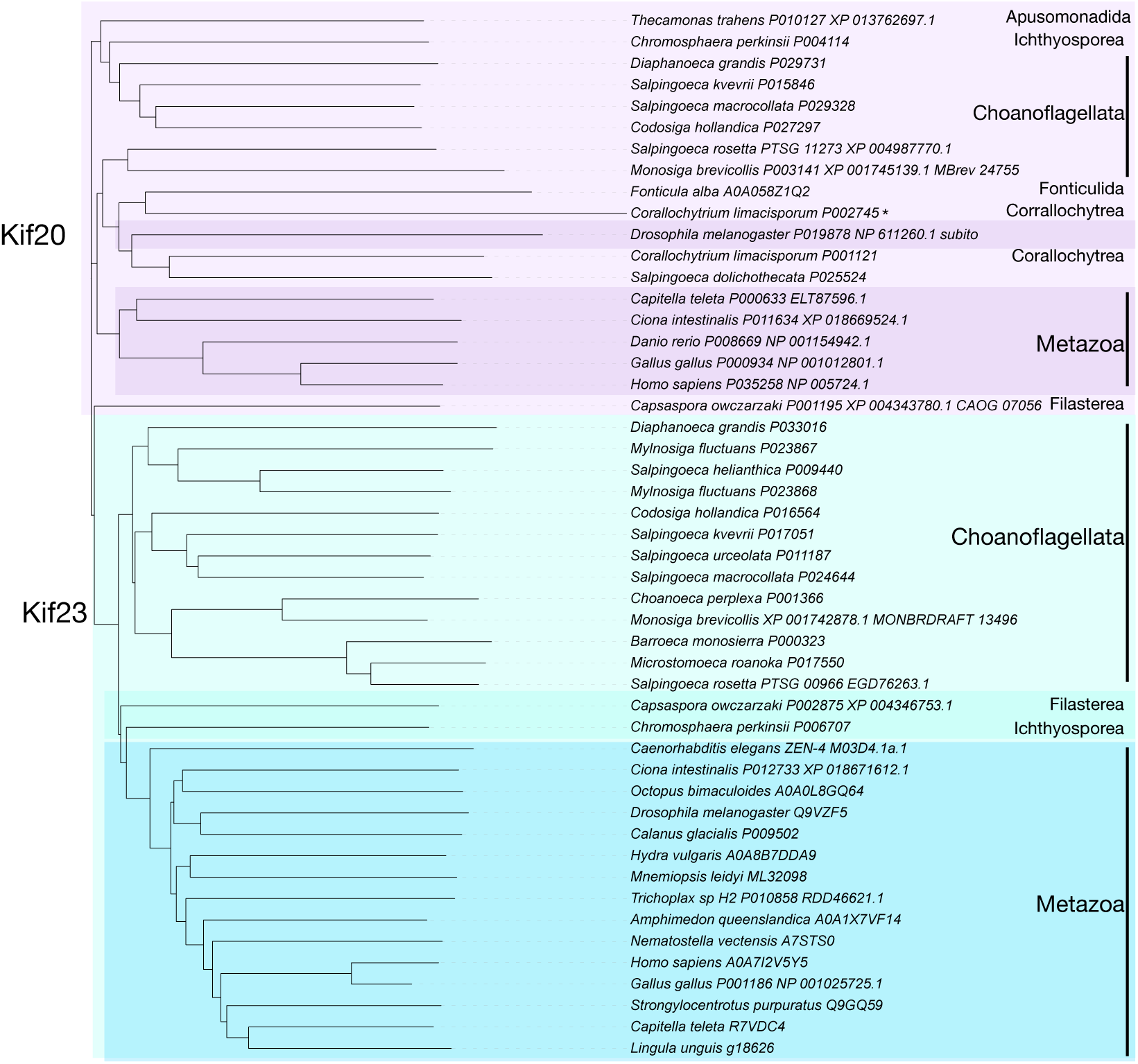
Phylogenetic tree of Kif20 and Kif23 in Metazoa, Choanoflagellata, and other Holozoa. Cluster alignment of kinesin-6 family members from Metazoa, Choanoflagellata, and other Holozoa. The motor domains of the indicated sequences were aligned using Clustal. Kif20 sequences are highlighted in shades of purple; Kif23 in shades of blue. Despite Corallochytrium limacisporum P002745 clustering with Kif20, it contains a predicted Arf6 binding domain (albeit low confidence).

**Supplementary Figure 2:**
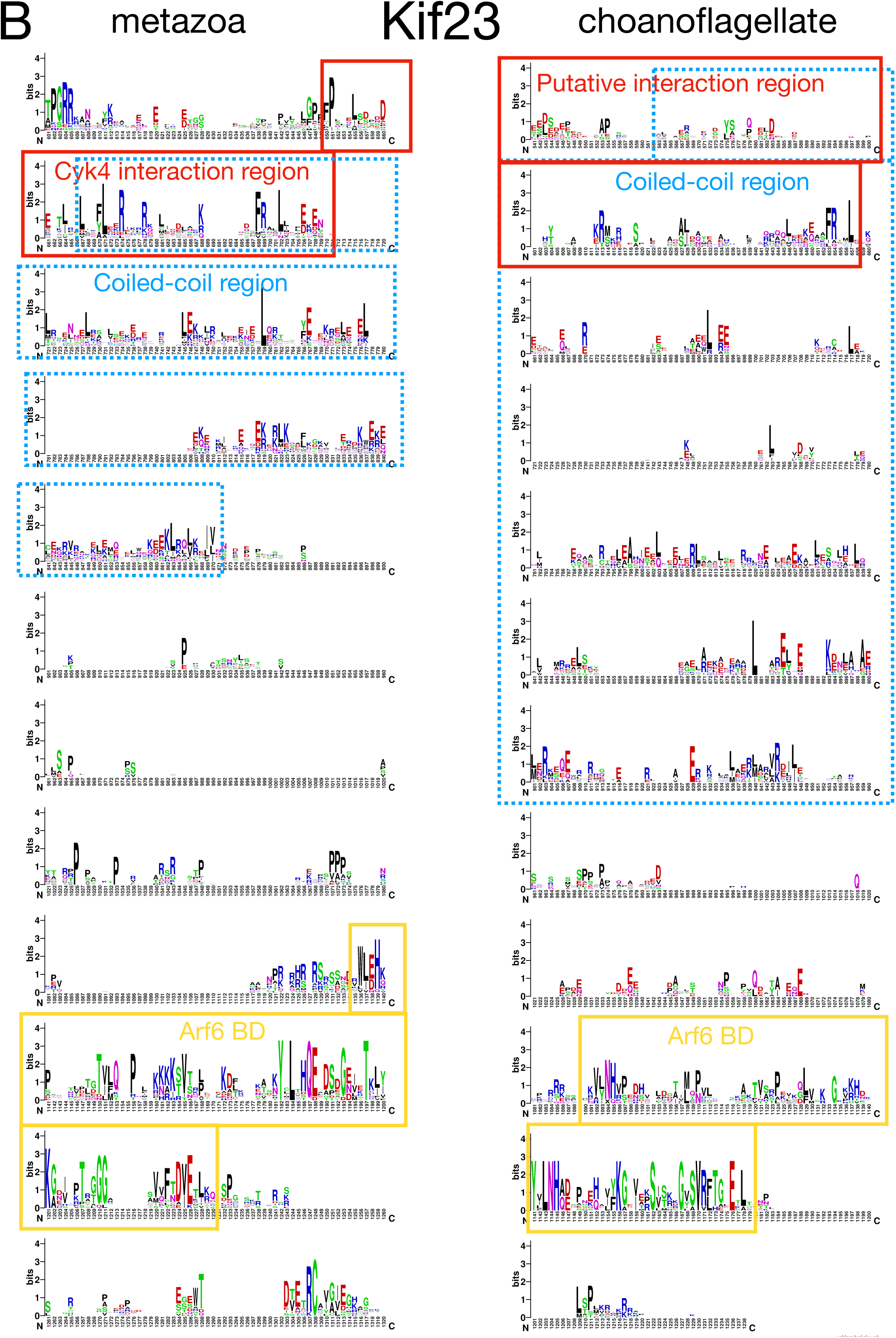
Sequence alignment of Kif23 from Metazoa and Choanoflagellata. Multiple sequence alignment of Kif23 family members from Metazoa and Choanoflagellata displayed using WebLogo. (A) Comparison of motor domains. (B) Comparison of region C-terminal to the motor domain. The interaction region is boxed in red, the coiled coil regions in blue, and the Arf6 binding domains in yellow.

**Supplementary Figure 3:**
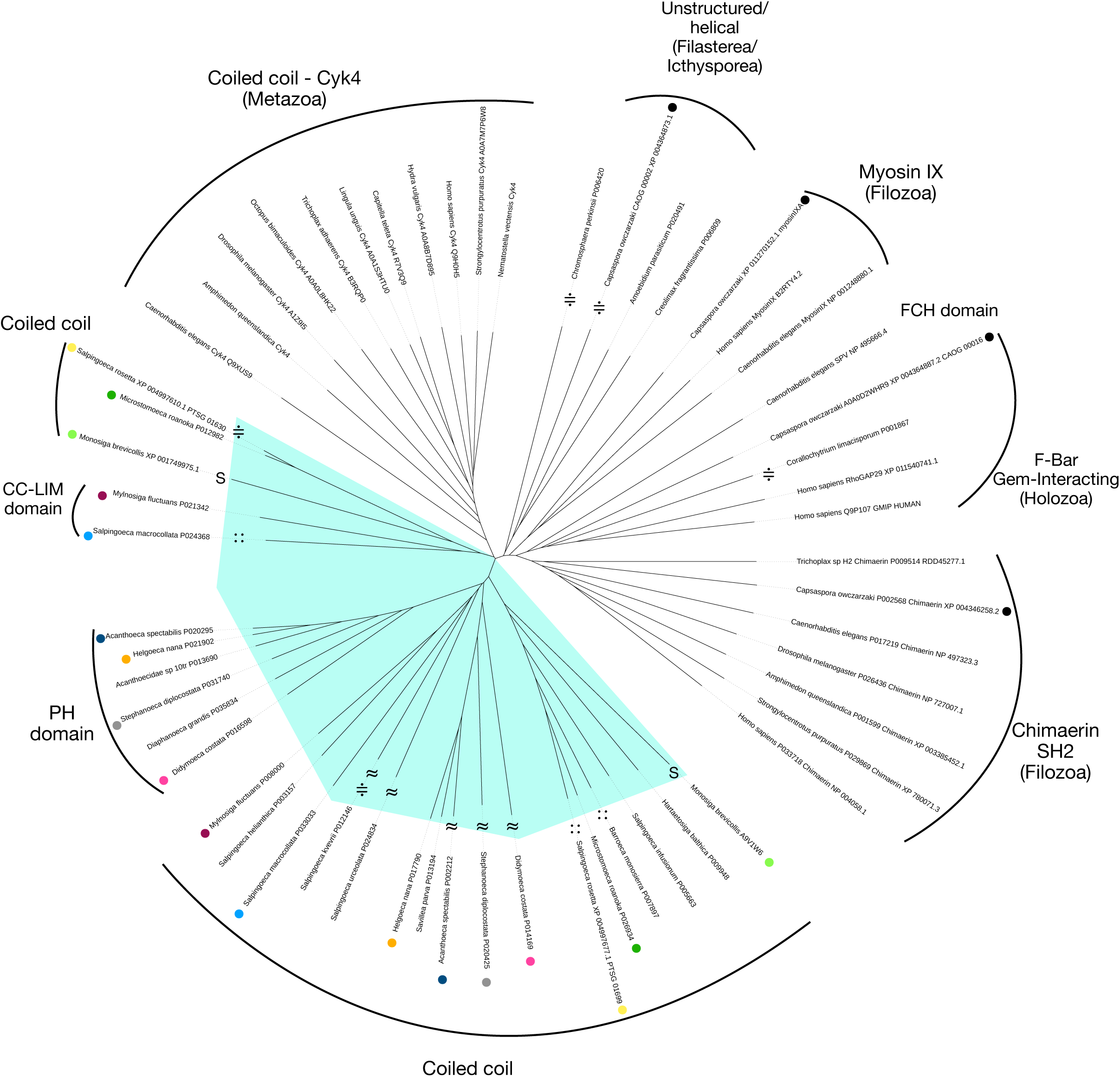
Phylogenetic tree of Cyk4 and Cyk4-related proteins in Metazoa, Choanoflagellata, and other Holozoa. Cluster alignment of selected C1:RhoGAP domain containing family members from Metazoa, Choanoflagellata (highlighted in turquoise), and other Holozoa. The C1:RhoGAP domains of the indicated sequences were aligned using Clustal and displayed as an unrooted tree. Paralogs from the same non-metazoan species are marked with a dot of the same color. The domain structure of the associated N-terminal domains are indicated. Proteins with a prediction of the N-terminal domain in figure 3 are marked “≈”, proteins with a prediction of the N-terminal domain and in complex with the cognate Kif23 in figure 3 are marked “∷”, proteins predicted to fail to form heterotetrameric complexes shown Supplementary Figure 6 are marked “≑”. Proteins marked with a (≈, ∷, or S) are included in Supplementary Figure 4.

**Supplementary Figure 4:**
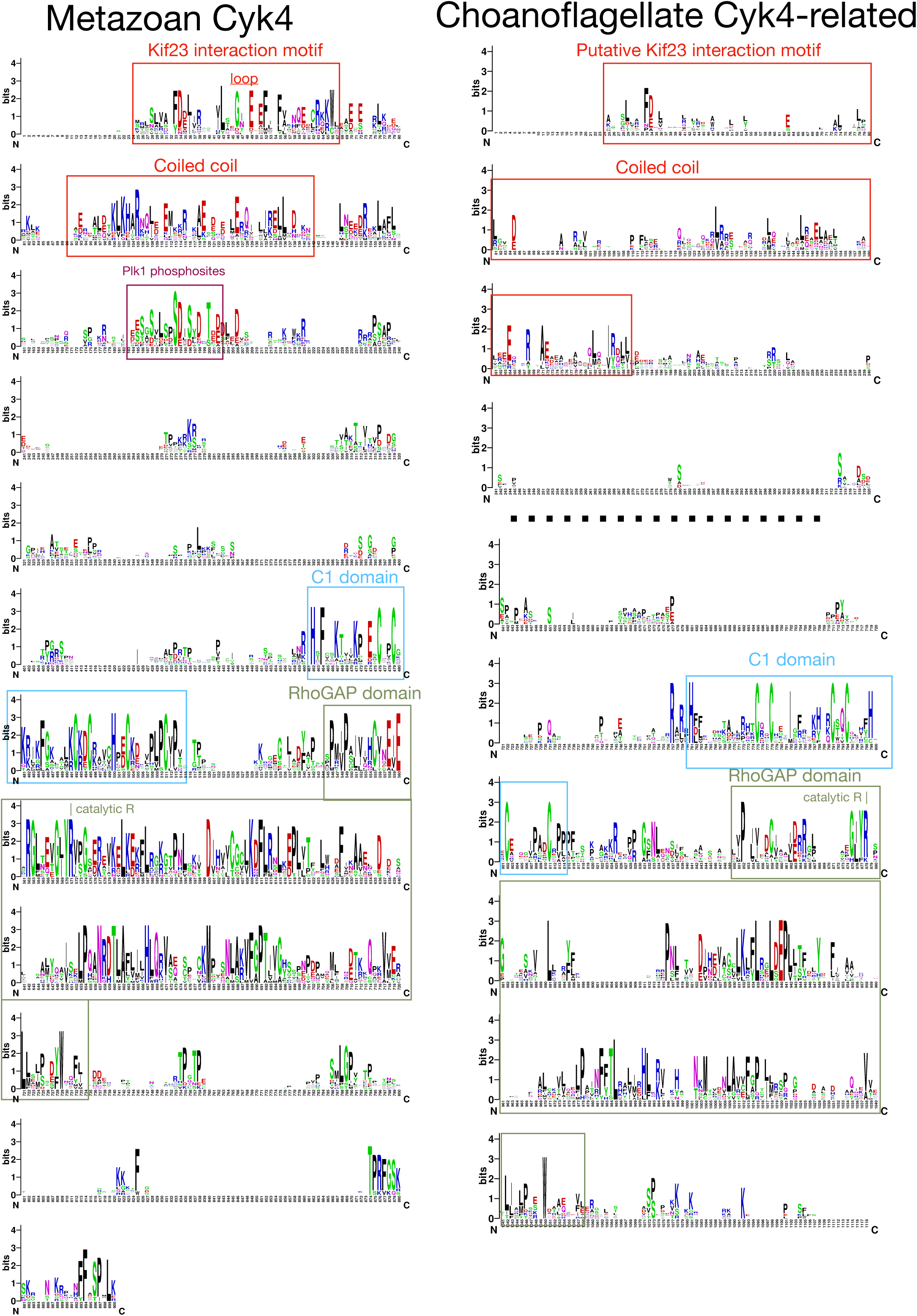
Sequence alignment of Cyk4 from Metazoa and Cyk4-related proteins from selected species of Choanoflagellata. Multiple sequence alignment of Cyk4 family members from Metazoa and Cyk4-related proteins from Choanoflagellata displayed using WebLogo. The metazoan alignment is constructed from all the metazoan Cyk4 proteins indicated in Supplementary Figure 3. The choanoflagellate alignment contains all the Cyk4-related proteins in Supplementary Figure 3 that are marked with the symbols (≈, ∷, or S). The interaction region and coiled coil are boxed in red, the C1 domain in blue, and the RhoGAP domain in olive green. A central region of low conservation of of the choanoflagellate Cyk4-related proteins was omitted (dotted line).

**Supplementary Figure 5:**
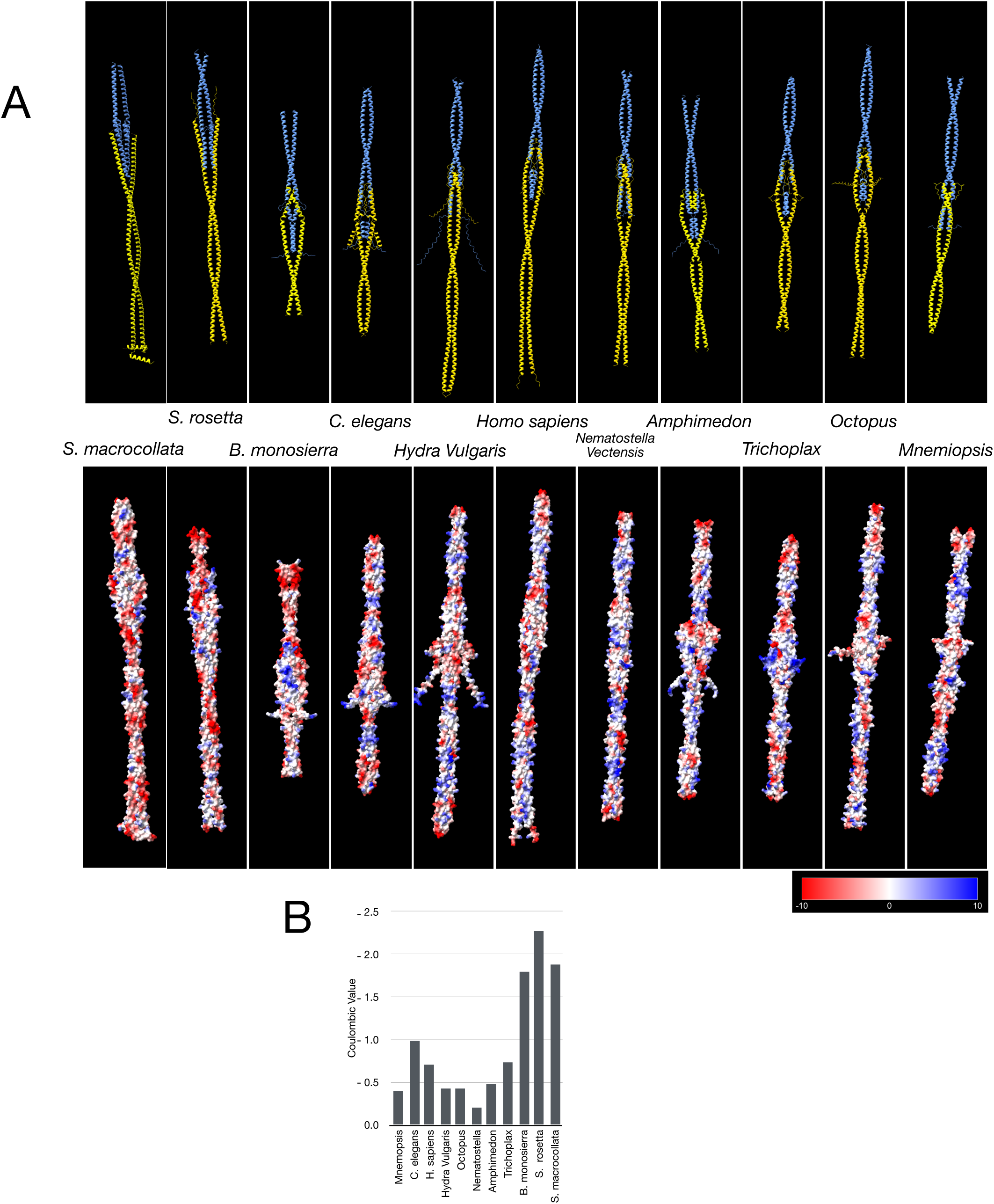
Visualization of the centralspindlin complex assembly region of Kif23 with metazoan Cyk4 or choanoflagellate Cyk4-related proteins. (A) Backbone structures of the predicted interaction regions of Cyk4 and Cyk4-related proteins (blue) and Kif23 orthologs (yellow). (B) Surface electrostatics of the complexes. (C) The mean surface electrostatic potential of each complex.

**Supplementary Figure 6:**
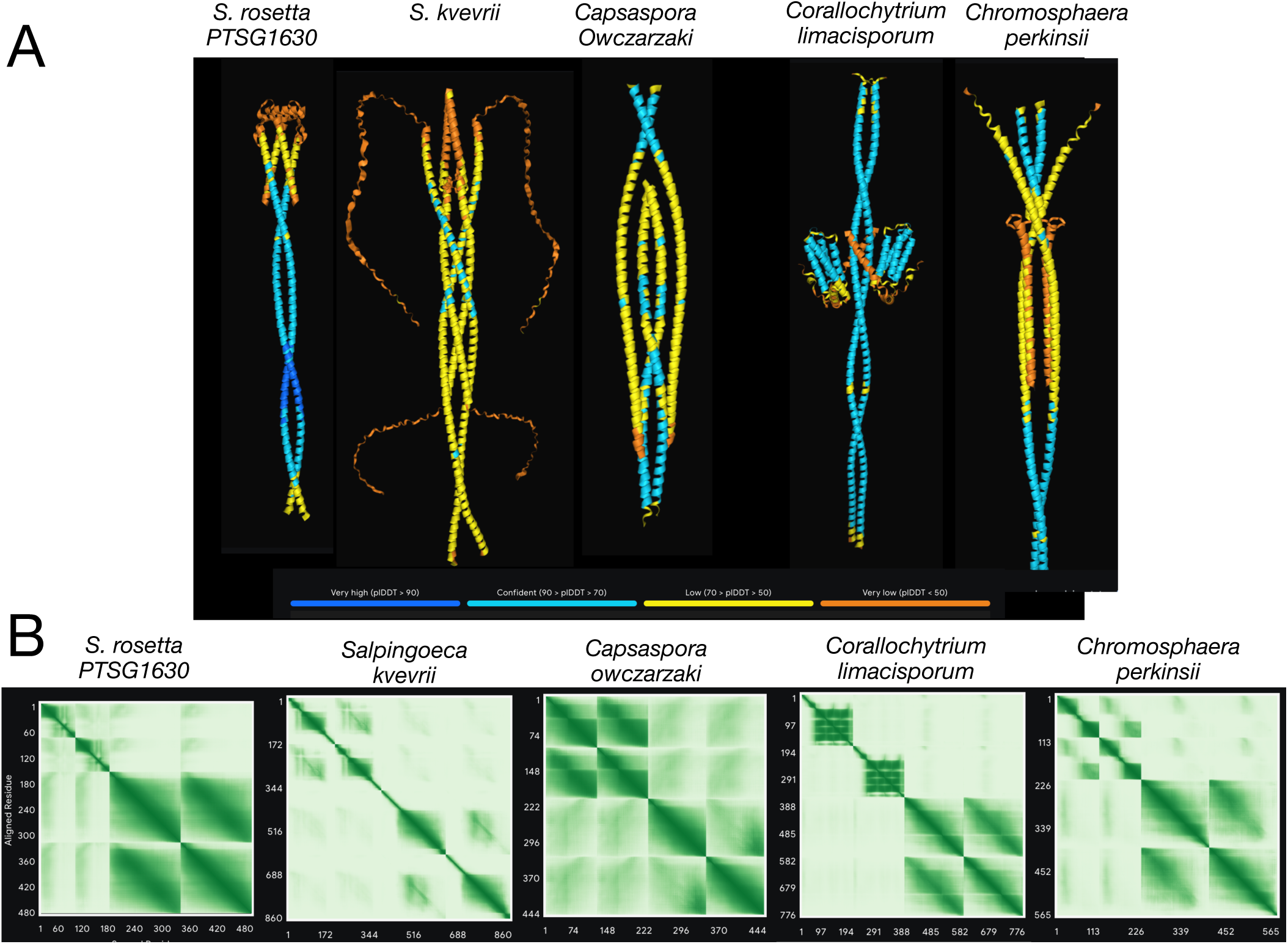
Visualization of putative assemblies of Kif23 and Cyk4-related proteins from Choanoflagellate, Filasterean, or Teretosporean species. (A) Alphafold3 predictions of potential complexes between Cyk4-related proteins and Kif23 from the indicated species with the associated PAE plots (B). The corresponding pTM and ipTM scores are displayed in figure 3E. The low confidence of the backbone predictions, the high interdimer PAE values, and the low pTM and ipTM scores suggests these proteins do not form stable complexes. Although the two subunits in the *Corallochytrium limacisporum* structure are each individually predicted with confidence, the Cyk4-related subunit surrounds the Kif23 coiled coil without evidence of a specific interaction.

