## Supplementary material for "Phylogenetics and structural modeling of centralspindlin and Ect2 provide mechanistic insights into the emergence of Metazoa and multicellularity": Sequence files: Gene predictions.pdf

This phylogenetic analysis depends upon gene predictions arising from various genome sequencing projects; many were labeled as “draft” or “preliminary” predictions. During the course of the analysis, some gene annotations appeared questionable and, where possible, revised. Such revisions are justified on the basis of parsimony and an assumption of protein conservation. These issues were particularly necessary for the analysis of species for which there is limited sequence coverage in the above databases (Ctenophora and Monosiga brevicollis). Sequence coverage of Placozoa and Porifera is also somewhat limited, as Kif23, Cyk4, and Ect2 were not all represented in a single species from these phyla, so representatives from other species were included, as indicated. All sequences are provided in the supplementary information.

```
Original
>MNELE06526 | ML05236a | [Mnemiopsis leidyi]
MMFLILSPLTPGQIGVRTTVIPQKYTSHSPIVSGSIQPSTRTGSRTGLRTGLRTGSRTGLAIPEPVREPVRTSYDMITKWIEFISNQADLQQRCHAAEGEVTRLRMQVNQY
KSDKEASDLRVKHIRKQLAEEKSQKQSLSKQNEKLIQQIRVIKEILEDRDRMSMMTDEDKTALGLNRSHLRKQFDKVFEDVSVDLDSDQSLEPSFEGAEELEETKTEWRRR
SRRSNVRKSDRMSTGQVAMKKRRTSNLHAPKEEMEPTTSESSEPPVIYPNLKKFNKANPVPVVAAEVISMSDATTISTTETESDAEYVQSDTESISSTTSESSLPNRAHHF
ALKSMIKPEGCKACRKRLGFSAHALRCKYCKMVVHVECKNAAPMPCVPVLSTSSKKQDGTLGSYVPKNPPHVPSILLRCVQEIEARGLNETGLYRVTGPDKAIKELKERML
RPRCQTNLRKVEDTTILTGVVKNFLMNLNDPILTSALSDEFKEYAEKGSTNHLYGTISKLPHENRHTLAFMIIHLQKVSESEHCKMPRSNIARVFAPTLVASIAMTDMDSA
MRDTKRSAKVVECLLDLPPSYWHNMLNPHDECSSPTPYSHDEAPHTPGTPEMLKGRNFFFLKLFLLTIKHSLTQNDIIDKMLLRSQCRSHLYSYEDCH
revised:
>MNELE06526_revised | ML05236a | [Mnemiopsis leidyi]
MSRKKSSSRQSLGAVRNTPSAYFDEICQISSILNCGLEQEWIEFISNQADLQQRCHAAEGEVTRLRMQVNQYKSDKEASDLRVKHIRKQLAEEKSQKQSLSKQNEKLIQQI
RVIKEILEDRDRMSMMTDEDKTALGLNRSHLRKQFDKVFEDVSVDLDSDQSLEPSFEGAEELEETKTEWRRRSRRSNVRKSDRMSTGQVAMKKRRTSNLHAPKEEMEPTTS
ESSEPPVIYPNLKKFNKANPVPVVAAEVISMSDATTISTTETESDAEYVQSDTESISSTTSESSLPNRAHHFALKSMIKPEGCKACRKRLGFSAHALRCKYCKMVVHVECK
NAAPMPCVPVLSTSSKKQDGTLGSYVPKNPPHVPSILLRCVQEIEARGLNETGLYRVTGPDKAIKELKERMLRPRCQTNLRKVEDTTILTGVVKNFLMNLNDPILTSALSD
EFKEYAEKGSTNHLYGTISKLPHENRHTLAFMIIHLQKVSESEHCKMPRSNIARVFAPTLVASIAMTDMDSAMRDTKRSAKVVECLLDLPPSYWHNMLNPHDECSSPTPYS
HDEAPHTPGTPEMLKGRNFFFLKLFLLTIKHSLTQNDIIDKMLLRSQCRSHLYSYEDCH

>XP_063677920.1 rac GTPase-activating protein 1-like [Bolinopsis microptera]
MTKKKSTSRQSLGAARNAPSAYFDELCQISTILNCGLEQEWIEFISNQADLQQRCQAAEGEVTRLRVQVN
QFKSDKEASDLRVKHMRKQLAEEKSQKQALSKLNEKLVQQIRVIKEILEDRDRMSMMTDEDKTALGLNKS
HLRKQFDKVFEDVSVDLDSDQSLEPSFEGELEETRTEWRRRSRRSNVRKSERQSTGPGVTVKKRRTSNLH
HPPKEDIYMTMEPTSSESSEPPVIYPNLKKFNKAHPVPHVEAEVISVQSSEATTISTTETESDVDYLPSD
TESISSTTSESSLPNRPHHFALKSMIKPEGCKACRKRLGFSAHALRCKYCKMVVHQDCKNAAPMPCVPIL
STPSRKQDGSLFSYVPKNPPHVPSIMLRCVQEIEARGLNEIGLYRVTGPDKAIKELKERMLRPRCQTNLR
KIDDTTILTGVLKNFLMNLNDPILTSALSDEFKNNAEKGSANHLYGTISKLPHENRHTLAFLIIHLQKVA
ESEHCKMPRSNIARVFAPTLVASIAMTDMDSAMRDTKRSAKVVECLLDLPPYYWTNMLNPHEDCSSPTPY
SNDEAPRTPGTPEMLKVQPGMYQPPTPTNAKETTFTQRVFGGKKKTFFNKFETP
```

CLUSTAL O(1.2.4) multiple sequence alignment

#### Alignment of draft Mnemiopsis gene prediction vs Bolinopsis

```
MNELE06526          MMFLILSPLTPGQIGVRTTVIPQKYTSHSPIVSGSIQPSTRTGSRTGLRTGLRTGSRTGL      60
XP_063677920.1      -----------------------------------MTKKKSTSRQSLGAA--RN-APSA      21
                                                          ..:: ** .* :.   .   .

MNELE06526          AIPEPVREPVRTSYDMITKWIEFISNQADLQQRCHAAEGEVTRLRMQVNQYKSDKEASDL      120
XP_063677920.1      YFDELCQISTILNCGLEQEWIEFISNQADLQQRCQAAEGEVTRLRVQVNQFKSDKEASDL      81
                     : *  :  .  . .:  :***************:**********:****:*********

MNELE06526          RVKHIRKQLAEEKSQKQSLSKQNEKLIQQIRVIKEILEDRDRMSMMTDEDKTALGLNRSH      180
XP_063677920.1      RVKHMRKQLAEEKSQKQALSKLNEKLVQQIRVIKEILEDRDRMSMMTDEDKTALGLNKSH      141
```

```
                          ****:************:*** ****:******************************:**

MNELE06526                LRKQFDKVFEDVSVDLDSDQSLEPSFEGAEELEETKTEWRRRSRRSNVRKSDRMSTGQ-V      239
XP_063677920.1            LRKQFDKVFEDVSVDLDSDQSLEPSFEG--ELEETRTEWRRRSRRSNVRKSERQSTGPGV      199
                          ****************************  *****:***************:* ***  *

MNELE06526                AMKKRRTSNLHAPKE-----EMEPTTSESSEPPVIYPNLKKFNKANPVPVVAAEVISM--      292
XP_063677920.1            TVKKRRTSNLHHPPKEDIYMTMEPTSSESSEPPVIYPNLKKFNKAHPVPHVEAEVISVQS      259
                          ::********* * :      ****:*******************:*** * *****:

MNELE06526                SDATTISTTETESDAEYVQSDTESISSTTSESSLPNRAHHFALKSMIKPEGCKACRKRLG      352
XP_063677920.1            SEATTISTTETESDVDYLPSDTESISSTTSESSLPNRPHHFALKSMIKPEGCKACRKRLG      319
                          *:************.:*: ****************** **********************

MNELE06526                FSAHALRCKYCKMVVHVECKNAAPMPCVPVLSTSSKKQDGTLGSYVPKNPPHVPSILLRC      412
XP_063677920.1            FSAHALRCKYCKMVVHQDCKNAAPMPCVPILSTPSRKQDGSLFSYVPKNPPHVPSIMLRC      379
                          **************** :***********:*** *:****:* *************:***

MNELE06526                VQEIEARGLNETGLYRVTGPDKAIKELKERMLRPRCQTNLRKVEDTTILTGVVKNFLMNL      472
XP_063677920.1            VQEIEARGLNEIGLYRVTGPDKAIKELKERMLRPRCQTNLRKIDDTTILTGVLKNFLMNL      439
                          *********** ******************************::********:*******

MNELE06526                NDPILTSALSDEFKEYAEKGSTNHLYGTISKLPHENRHTLAFMIIHLQKVSESEHCKMPR      532
XP_063677920.1            NDPILTSALSDEFKNNAEKGSANHLYGTISKLPHENRHTLAFLIIHLQKVAESEHCKMPR      499
                          **************: *****:********************:*******:*********

MNELE06526                SNIARVFAPTLVASIAMTDMDSAMRDTKRSAKVVECLLDLPPSYWHNMLNPHDECSSPTP      592
XP_063677920.1            SNIARVFAPTLVASIAMTDMDSAMRDTKRSAKVVECLLDLPPYYWTNMLNPHEDCSSPTP      559
                          ****************************************** ** ******::******

MNELE06526                YSHDEAPHTPGTPEMLKGRNFFFLKLFLLTIKHSLTQNDIIDKMLLRSQCRSHLYSYEDC      652
XP_063677920.1            YSNDEAPRTPGTPEMLKVQPGMYQPPTPTNAKETTFT-----QRVFGGKKKTFFNKFETP      614
                          **:****:********* :  ::      . *.:        : :: .: ::.: .:*

MNELE06526                H     653
XP_063677920.1            -     614
```

### Alignment of revised Mnemiopsis gene prediction vs Bolinopsis

```
XP_063677920.1            MTKKKSTSRQSLGAARNAPSAYFDELCQISTILNCGLEQEWIEFISNQADLQQRCQAAEG 60
MNELE06526_revised2       MSRKKSSSRQSLGAVRNTPSAYFDEICQISSILNCGLEQEWIEFISNQADLQQRCHAAEG 60
                          *::***:*******.**:*******:****:************************:****

XP_063677920.1            EVTRLRVQVNQFKSDKEASDLRVKHMRKQLAEEKSQKQALSKLNEKLVQQIRVIKEILED 120
MNELE06526_revised2       EVTRLRMQVNQYKSDKEASDLRVKHIRKQLAEEKSQKQSLSKQNEKLIQQIRVIKEILED 120
                          ******:****:*************:************:*** ****:************

XP_063677920.1            RDRMSMMTDEDKTALGLNKSHLRKQFDKVFEDVSVDLDSDQSLEPSFEG--ELEETRTEW 178
MNELE06526_revised2       RDRMSMMTDEDKTALGLNRSHLRKQFDKVFEDVSVDLDSDQSLEPSFEGAEELEETKTEW 180
                          ******************:******************************  *****:***

XP_063677920.1            RRRSRRSNVRKSERQSTGPGVTVKKRRTSNLHHPPKEDIYMTMEPTSSESSEPPVIYPNL 238
MNELE06526_revised2       RRRSRRSNVRKSDRMSTGQ-VAMKKRRTSNLHAPKE-----EMEPTTSESSEPPVIYPNL 234
                          ************:* ***  *::********* * :      ****:*************

XP_063677920.1            KKFNKAHPVPHVEAEVISVQSSEATTISTTETESDVDYLPSDTESISSTTSESSLPNRPH 298
MNELE06526_revised2       KKFNKANPVPVVAAEVISM--SDATTISTTETESDAEYVQSDTESISSTTSESSLPNRAH 292
                          ******:*** * *****:  *:************.:*: ****************** *

XP_063677920.1            HFALKSMIKPEGCKACRKRLGFSAHALRCKYCKMVVHQDCKNAAPMPCVPILSTPSRKQD 358
MNELE06526_revised2       HFALKSMIKPEGCKACRKRLGFSAHALRCKYCKMVVHVECKNAAPMPCVPVLSTSSKKQD 352
                          ************************************* :***********:*** *:***

XP_063677920.1            GSLFSYVPKNPPHVPSIMLRCVQEIEARGLNEIGLYRVTGPDKAIKELKERMLRPRCQTN 418
```

```
MNELE06526_revised2      GTLGSYVPKNPPHVPSILLRCVQEIEARGLNETGLYRVTGPDKAIKELKERMLRPRCQTN 412
                         *:* *************:************** ***************************

XP_063677920.1           LRKIDDTTILTGVLKNFLMNLNDPILTSALSDEFKNNAEKGSANHLYGTISKLPHENRHT 478
MNELE06526_revised2      LRKVEDTTILTGVVKNFLMNLNDPILTSALSDEFKEYAEKGSTNHLYGTISKLPHENRHT 472
                         ***::********:*********************: *****:*****************

XP_063677920.1           LAFLIIHLQKVAESEHCKMPRSNIARVFAPTLVASIAMTDMDSAMRDTKRSAKVVECLLD 538
MNELE06526_revised2      LAFMIIHLQKVSESEHCKMPRSNIARVFAPTLVASIAMTDMDSAMRDTKRSAKVVECLLD 532
                         ***:*******:************************************************

XP_063677920.1           LPPYYWTNMLNPHEDCSSPTPYSNDEAPRTPGTPEMLKVQPGMYQPPTPTNAKETTFTQR 598
MNELE06526_revised2      LPPSYWHNMLNPHDECSSPTPYSHDEAPHTPGTPEMLKVQPGMYQPPTPTNSKETTFTQR 592
                         *** ** ******::********:****:**********************:********

XP_063677920.1           VFGGKKKTFFNKFETP  614
MNELE06526_revised2      VFGGKKKTFFNKFETP  608
                         ****************
```

Mnemopsis Ect2 ortholog

The closest annotated ortholog to Ect2 in Mnemiopsis is ML444210a:

```
>ML444210a [from Mnemnopsis genome project]
MYHDRPEDSEMNDFITQLGIENPATLDKSEIGSAVRQYAVLPDFEMGEKFLSKLQARHVRVISPVLIKQCLVSGRPLGSFRRFLMPGRRCYPMFSSMMVGLTIGFTNFPSR
EDLKRMRWMVEVLGGRVTREYDPMGKIILIANMAGRDKFETAMACGEPVLKVDWLNDCYSLRMEPNVLPRLMYAEDNQRYACEIFYRFKFFLIGFKEYDAVDIKENCKRYG
AVVFEEVSVMRKCTHVIIYESSDQSAYQPYLEELRKICSREGDLSTQLVTDKWFYESVSLSALLDTNRYSITISPDGTLDFSSAHDRKRRVDELSADSCPRKKRSSEANCS
IYSNISITGQLPKTPKNTPKTIDKRDNVISEFIRTEETYVRHLKIIEERYYGYLYEQVANPPANNKFELELPSVHMMFQKIVPIRIVHGALLKELERYEVSEKLRKDSKKN
RGEKSFGRILLNWIGKEVFKDENAPDGISFLNYSREEMIKMFQYDKNETRDYLRFDPGRTSSEKRDEGERRRDQEKRRKEYEGTMRKKPSGEVQGEDPDVLKHYNIYMNAH
EGILDLINKNETKCFSEKIEKKLLTLLVKYWVSELQLLVFCNFSLCRERTVTVQENVRQTSNTPGHSRKSGVGTKVYPLGGNVDFCPLRRGSCMCQGAGSELLELPPRTLL
RSGRCTRTCRHSPADTPSPGPRSSSRSRHLGEKSGCRGASIPRT
```

This gene model is likely to be partially mis-annotated. The first ~484 amino acids are highly homologous to Ect2 (the top hit vs C. elegans is ECT-2), but the sequences diverge at that point. The first 270 amino acids are predicted to form ~2+ tandem BRCT domains. However, the DH domain is incomplete and the PH domain is absent. The transcriptomic databases of the  Mnemiopsis genome project contains additional predictions (ML4442.24008.3, ML4442|comp13217_c0_seq1, MLRB444219) enabled the reconstruction of a full length Ect2 with all domains intact.

Monosiga brevicollis Cyk4 paralogs

The Monosiga brevicollis protein MONBRDRAFT_34379 (https://www.ncbi.nlm.nih.gov/gene?cmd=retrieve&list_uids=5895264 ; https://www.ncbi.nlm.nih.gov/protein/XP_001749975.1) derived from the genome sequencing project {King et al., 2008, #178407} contains a C1:RhoGAP domain. The first two exons of this predicted protein is replete with repeats of the dipeptides TH (12) and TE (47). The first ATG of this protein is more likely to be at position 188722; this prediction was used for further analysis.

*Monosiga brevicollis* A9V1W6 (aka XM_001746567.1) is predicted to encode a 1365 aa protein containing a RhoGAP domain at 436-636. Following the RhoGAP domain, in the genomic sequence corresponding to  amino acid 657, there is a predicted intron of 530 bp, longer that the average intron size of in this organism of 174 nt. The protein sequence that follows in exon 9 and beyond (listed below) is 35% identical to bacterial proteins containing a DUF6259 domain (e.g. https://www.ebi.ac.uk/interpro/entry/InterPro/IPR046226/alphafold/#table). Based on the protein size, intron length, and unusual homology this draft gene annotation is likely erroneous. Here, the analysis of the protein was limited to the N-terminal RhoGAP domain containing portion.

```
>C-terminus of A9V1W6
KKQNISLLLTLTTPSSLLVLNTDGSIADLRARTANPGNWSFVLDAHRPIFTFLASGVEPTGPTSNISARDCKAWPPELVNATQARVRFLCPSTLVPA
```

```
AAFDITVSVQADQERDQFLFGLELNINATNASVVLSALEFPLLTMAAIRDEMLLSEDNIVMPYAGGTRNNMPGIASEAQSQMYPGYASFQFWARSGF
KALKPHRFVILHVKEFVYTSAAHDSINITVRHHRPWIPFDAATPFLPYQTALRTFSGDWMTAATLYKAWAGQQWWTRTPLTRRTDIPSFLLNGTGVW
ISGINSQSGFNPNRWEHDGTWAGIDYLPPQPSADAWQNATQQLLAQSSNGPAFNHTTQMLAHQSMLVRNNDQTLAAVDAYNEPEGPQGWRGYSTRLC
HGSREAQTVLGAIFHQVSALNVSLISFDQEIGGGARYPCYANNHGHALGWGPWMYHEFAQLVQEIKANATHQGIMTEQVSELTIPLMATYWSRQFAQ
QHYFFFRGYGEPIFNFLYHEFVTTIGAAEVQGQGEVAQYRSVVLRRAVIGDCVARGMLQGPFDSDVSLAPVPGSFSQNISMLFFNISATYSHYAEYV
TLGQMVRPPISRNPVDLTTTLVWKRGNGSFVREAVNTSALRLGAFMAPEGTVGILIVNIDDVTHSVTLDITQTWKPPPNATVMIYDEQCRPMGTAKV
EQSGSTVQLVVPGLRLRFLALSPPSCAQ
```

*Monosiga brevicollis* Kif23 paralog

One relevant gene from *Monosiga brevicollis* is MONBRDRAFT_13496, (XP_001742878.1). This kinesin is the closest ortholog of *S. Rosetta* kinesin PTSG00966, but it is a partial sequence. The annotation specifies that it is N-terminally truncated. The predicted protein is missing the initiator methionine, but more importantly for these purposes it is missing the parts after the motor domain, which would be the region that could interact with Cyk4.This truncated protein was used in the analysis to the limited extent possible, with the understanding that it is undoubtedly incomplete.

```
>XP_001742878.1 uncharacterized protein MONBRDRAFT_13496, partial [Monosiga brevicollis MX1]
CSLQQVDIEHDDVRYGVFVQYVEIYNNYCYDLLVPPPPEKEPAKPRPPTRQSLKISQDETGHRYVREAIWKEVHTPEEAFELLHQGQITRATAATDL
NATSSRSHSIFTIRLVSAPTSMRGNDVQRDGPTPHVADISLVDLAGSERTSRTNSTGDRRKEAGAINKSLSVLRDCMKALRRNQDNGSRGKPKFNES
SLTKLFERFLTGDGLAAMVVCASPAQSDASETAHVFDFAKVASDVQVEAPVHVMQDHGKLLDHGVRTHTRPACQEQCKPRTFGSFITRLVSFIEWKS
```
